# Comparative phytotoxicity and chironomid toxicity assessments of leaf successive extract fractions of Asiatic poison bulb, *Crinum asiaticum* L

**DOI:** 10.1101/2020.12.15.422968

**Authors:** Souren Goswami, Sanjib Ray

## Abstract

*Crinum asiaticum* is an evergreen bulbous perennial shrub of Amaryllidaceae family with ethnomedicinal importance and our earlier study described a comparative account antimicrobial and antioxidant properties of the different solvent-mediated sequential extract fractions. The present study aimed to analyze their comparative account of induced phytotoxicity and chironomid toxicity. For phytotoxicity assessment, germination inhibition and seedling’s root and shoot growth retardation effects on *Triticum aestivum and Cicer arietinum* were analyzed and for lethal concentration determination, the freshwater bottom-dwelling first instars chironomid larvae were used. The crude aqueous, petroleum ether and ethyl acetate extracts showed significant toxic effects on both meristematic tissue and aquatic midges. The phytotoxic assays indicate that the ethyl acetate fraction contains the most potent growth inhibitors, followed by the crude aqueous and petroleum ether fractions. The last aqueous fraction was found to be the least toxic, the highest LC_50_ and LT_50_ values and ethyl acetate extract fraction having highest toxicity. Thus the present study supplements to our earlier report, that indicated the last aqueous extract fraction of *C. asiaticum* has potent antioxidant and antibacterial potentials as well as its prospective use in livestock maintenance, as it is least toxic and the ethyl acetate extract, the most toxic fraction identified here, is needed to explore for pharmaceutical importance.

## Introduction

The Ethno-medicinal plants and their differential uses in folk cultures have a vital role in drug discovery. Even today, plants are used as a source of remedies for several fatal diseases when allopathic treatment fails. The herbal remedies remain as an ever popular in the health care in different populations in different names. In drug discovery research, a massive number of plants are currently being explored for active principles. Recently we have explored a comparative account of antioxidant and antimicrobial potentials of the different extract fractions isolated from the leaves of *Crinum asiaticum* (Goswami et al., 2020).

*Crinum asiaticum* (Commonly: Asiatic Poison Lily) a tuberous herb of Amaryllidaceae family. This plant is distributed as ornamental throughout tropical India and is native to tropical south-eastern Asia (China, Hongkong, Srilanka, Myanmar, Thailand, Malaysia, and Japan) mostly on sandy seacoasts and the back mangroves (Ghani, 1998; Zhanhe and Alan, 2000; Walter, 1998; Davison *et al*., 2008). It is an evergreen small bulbous perennial shrub producing a rosette of fleshy glabrous, glossy leaves growing to a height of 1.5 - 2.0 m tall with a pointed tip, arises from an underground fleshy bulb that can be 5-15 cm in diameter. The greenish leaves of the plant are narrowly lanceolate, 1 −1.5 m long and 5 - 7 cm wide and appear as if sprouting from the base or ground (Kumar, 2011). The flowers appear in a cluster of white, sweetly fragranced, showy flowers and emerge from the middle of the trunk and are seen from the month of April-August (Hutchinson, 1964; Patel ei al., 2010). The flowering stalk is about 1 - 1.2 m long. The fruits are a little and irregularly rounded dehiscent capsule (about 5 cm) containing green seeds.

All parts of this plant are considered both poisonous and ethnomedicinally important. The leaves are used by Malays to treat several health complains including rheumatic remedies (Hutchings *et al*., 1996). A preparation from leaves is used to treat hemorrhoids, piles, inflamed joints, injury or abscesses and fractures (Wee, 1992; Singh *et al*., 2010). In Congo, it is used to treat leprosy. In Bangladesh it has enormous tribal use starting from simple pain-infestations to arthritis-leprosy (Ghani, 1998; Zhanhe and Alan, 2000; Walter, 1998).

Antimicrobial potential reported against some human pathogenic bacteria concluded it as a natural source of antibacterial compounds (Goswami et al., 2020; Ilavenil *et al*., 2010; Paul *et al*., 1995). A mannose-binding lectin (agglutinin) gene, cloned from this plant shows a great similarity to gastrodianin type antifungal proteins. Leaves extract demonstrated cytotoxic activity against murine P388 D1 cells (Ahmad, 1996). Lycorine present in hot water extract of *C. asiaticum* showed strong cytotoxicity against four human tumor cell lines i.e. A549, HL-60, LOVO and 6TCEM (Sun *et al*., 2009). Lycoriside, an alkaloid from *C. asiaticum* elicited concentration-dependent anti/pre-release effect on mast cell mediators of albino rats (Ghosal *et al*., 1985). Galanthamine, from similar origin, is an approved remedial for Alzheimer’s disease (Kogure *et al*., 2011). Two independent studies demonstrated, its ethanolic extract’s antioxidant nature, having protective role against oxidative damage in hepatic tissues of diabetic rats and human erythrocytes (Ahmad *et al*., 2011; Ilavenil *et al*., 2011). *In vivo* study of its leaf extract on both mice and rat models supports its traditional use as an analgesic agent and in the same model; the chloroform fraction of methanol extract significantly reduced the carrageenan-induced acute edema, supporting the previous claim (Samud *et al*., 1999; Asmawi *et al*., 2011; Rahman *et al*., 2011). Crinumin, a glycosylated serine protease, from the latex of *C. asiaticum*, shows plasmin-like thrombolytic activity and thus inhibits platelet aggregation and related diseases could be prevented and treated efficiently by crinumin. This herb remarkably decreases the lead content of soil as a phytoremedial tool to reduce heavy metal pollution. The plant exhibits all the major chemical traits of the Amaryllidaceae family such as alkaloids, flavonoids, coumarins, terpenoids, amides and phenolics (Fennell and van Staden, 2001; Sun *et al*., 2009, Goswami and Ray, 2017). 21 alkaloids have been characterized from it, with significant structural and pharmacological diversities (Kogure *et al*., 2011). Those alkaloids identified could be classified into four types i.e., Galanthan type, Galanthamine type, β-Ethano Crinan Type and α-Ethano Crinan Type. Among other major chemicals another 2 new alkaloids, asiaticumines A and B; 4 amides; 3 flavonoids; and 5 phenolics were also reported (Sun *et al*., 2009).

From the above-cited literature and our screening results, it is clear that *C. asiaticum* is a rich source of phytochemicals and with ethnomedicinal value; however, comparative account of toxicity of aerial parts’ extracts of *C. asiaticum* are not well studied. Therefore, this study aimed to investigate the comparative toxicity assessment of the aerial parts’ aqueous and the successive organic solvent mediated extract fractions of *C. asiaticum* using root apical meristem cells and Chironomid larvae as experimental model systems.

## 2. Materials and methods

### 2.1. Chemicals

Methanol and glacial acetic acid were obtained from BDH Chemicals Ltd., UK. Ascorbic acid was obtained from Merck Ltd., Mumbai, India. All other chemicals used in this study were of analytical grade and were obtained from reputed manufacturers. Silica gel of SRL, Mumbai, India

### 2.2. Plant collection, processing, and storage

Fresh aerial (leaves) parts of *C. asiaticum* were collected in a large scale from The Burdwan University campus, Golapbag, Purba Bardhaman, West Bengal, India. This plant species was taxonomically identified by Prof. Ambarish Mukherjee (Taxonomist), The Department of Botany, The University of Burdwan. A voucher specimen is maintained in the department for future reference. The collected plant materials were cleaned properly under tap water; shade dried; crushed into small pieces and pulverized using an electric grinder. The ground leaf powder was stored in an airtight glass container for future uses.

### 2.3. Extraction through Soxhlet apparatus

Aqueous extract (CaLAE) was prepared according to the standard procedure as described in our previous publication (Goswami and Ray, 2017), briefly, 50 g of leaf powder, extracted in 1 L distilled water for 24 h at 60°C, then filtered, condensed and stored at −20°C. For further detailed and comparative study different organic solvents were used in addition to distilled water to make the extracts. In a Soxhlet apparatus, 20 g dried leaf powder was extracted in each cycle, sequentially with 0.5 L of different solvents having increasing polarity with controlled temperature, i.e. 5 to 10°C beneath the steaming point of the respective solvent. Petroleum ether (PE), chloroform (Chl), ethyl acetate (EA), methanol (Me) and distilled water (Aq), were used sequentially to make the extraction complete; with minimum 72 h in each solvent and abbreviated respectively as CaPE, CaChl, CaEA, CaME and CaAq. Extracts were filtered through No. 1 Whatman filter paper (GE Healthcare UK Limited, Buckinghamshire, UK) and concentrated in a vacuum hot air oven, at a temperature lower than extraction, the final volume was recorded, and stored in −20°C. To calculate the concentration and extract value, in the hot (60°C) air oven, in five separate Petri dishes 5 mL extract (total 25 mL) was evaporated to complete dryness.

### 2.4. Experimental models

To study toxicity beside animal models both monocot (wheat) and dicot (chickpea) plant models were also used. Wheat (*Triticum aestivum*) seeds were used as a model to test germination inhibitory effects of the extracts. Wheat and chickpea (*Cicer arietinum*) seedling’s root and shoot growth inhibition were analyzed for phytotoxic and allelopathic activity assessments.

Larvae of the widespread dipteran, chironomids are the most abundant group in aquatic habtitats (Armitage et al., 1995; Pinder LCV, 1986; Benke AC, 1998). Their life cycle comprises of four larval instars and a pupal stage to reach the adult flying midge from the egg mass (Armitage et al., 1995; Pinder LCV, 1986). Overall, this frequent macro-invertebrates also serves a valuable food resource for the higher tropic levels, starting from insects-crustaceans to higher vertebrates and as well as mammals. These freshwater bottom-dwelling larvae have been used frequently in long-term laboratory toxicity assays. Agrochemical industries have been performing acute chironomid mortality (immobilization) tests for a number of years, thus, this is an ideal method of toxicity assay of chemicals. The test organisms were not maintained continuously rather collected freshly from local water bodies and maintained in the same water resource, kipping within a climate controlled chamber at 20±2°C with a 16:8 light/dark regime.

### 2.5. Germination inhibition assay

Surface sterilized (using 1 % sodium hypochlorite) healthy wheat seeds were allowed to germinate in a BOD under controlled condition (25±2°C), on sterilized wet filter paper in autoclaved glass Petri dishes, containing different test extracts in 1 mg/mL concentration. Petri dishes, having distilled water were considered as the control set. 35 seeds/Petri dishes and 5 replicas for each test samples (thus total 175 seeds/samples) were maintained for 3 days with a single treatment at the very beginning and in every 24 h required amount of distilled water was added and mixed properly to maintain the moist condition. Data were accumulated at 72 h of the experimental setup and from that germination percentage and its inhibition percentage were calculated.

### 2.6. Monocot and dicot growth retardation assay

Bioassays for growth retardation effects of the six test extracts (crude aqueous and five other soxhlet fractions) were done using both monocot (*T. aestivum*) and dicot (*C. arietinum*) seeds as experimental models. In monocot, the treatment dose was 1 mg/mL but in dicot, it was both 1 and 2 mg/mL. Seeds were germinated in distilled water after surface sterilization. For wheat seedlings, after 36 h of incubation 12 seeds of similar size were taken for each set and treated with the extracts. At the time of treatment (00 h), the root-shoot length was 00 mm and it was recorded up to 7 days in every 24 h intervals and the root-shoot growth inhibition percentages were also calculated. In dicot, ten germinated chickpea seeds of the parallel stage were taken for each set, and root length was measured at day 1^st^, 2^nd^, 3^rd^, and 6^th^. And from this growth inhibition was calculated as percentage values. In monocot after seven days of incubation accumulated dry weight of seedlings was measured. Without seeds, only root-shoot part was collected and dried in a hot air oven at 50°C for 48 h then weight was measured and compared with the control set maintained in distilled water.

### 2.7. Larvicidal assay of CaLAE and successive solvent fractions of C. asiaticum

Chironomid larvae are generally used as the standard aquatic test organism, representing the aquatic fauna, in the environmental risk assessment of any toxic substances (Weltje L, et al. 2010; OECD/OCDE 2011). As mortality is very hard to find in the larvae, conventionally immobility is used to next to mortality. In this study, acute toxicity is measured with first instars larvae of *Chironomus sp*., who were exposed to different range of concentrations of the test extracts in water-based medium for a period up to 96 h according to the protocol of Japanese chironomid acute toxicity guideline (JMAFF, 2005) and generic test guideline for mosquito larvae (WHO, 2005). After producing a gentle stream with a pipette, immobilization/mortality was recorded at 3, 24, 48, 72 and 96 h, and from this the median lethal concentration (LC_50_) values i.e. where 50% of the individuals are immobilized, at 24 h were determined for each test sample. As free-swimming first instars larvae are not stressed by the sediments and represent the most sensitive larval stage, they were used in this assay (Weltje L, et al. 2010; Treverrow N, 1985; Gauss et al., 1985). The stock solution and diluted test solutions were made with the pond water, of which the larvae belong to. And control was maintained in the same water without any addition or dilution. 30 larvae were exposed divided over 3 replicates of each test concentration for up to 96 h. Mortality/immobility percentages were counted and plotted against test concentrations. Data were analyzed to determine the concentration-response curves and the LT_50_ values of 1 mg/mL concentration i.e. time (in hours) required for killing off half of the individuals, with minimum 95% confidence limits (Stephan, 1977; Finney, 1978) were determined.

### 2.8. Scoring and Statistical analysis

Seed germination and seedling growth retardation percentages were calculated from triplicate sets. The obtained data set were analyzed using Microsoft Office Excel 2007 and Origin 8.0 software. Root-shoot lengths were expressed as Mean±SEM. Variations of data between treated and untreated groups were analyzed with the Student’s t-test and were considered statistically significant at least below *p*<0.05 level.

## 3. Results

### Screening of the most toxic fraction of C. asiaticum

### 3.1. Germination inhibition assay on wheat

The crude aqueous extract and other five successive organic soxhlet fractions of *C. asiaticum* leaf were tested on wheat seedlings to study their germination inhibitory effects. At 72 h of incubation, CaEA was found to be the most effective germinator inhibitor, followed by CaLAE, CaPE, CaAq, and CaChl (figure 01). They all significantly inhibited germination at the level of 0.1%. The CaMe showed no significant effect on wheat germination. The CaEA, CaLAE, and CaPE have inhibited germination more than half, on average, respectively as 81.1, 69.52 and 50.62% in respect to the control group. The last fraction (CaAq) inhibited only 11.6% germination.

**Figure 01.**
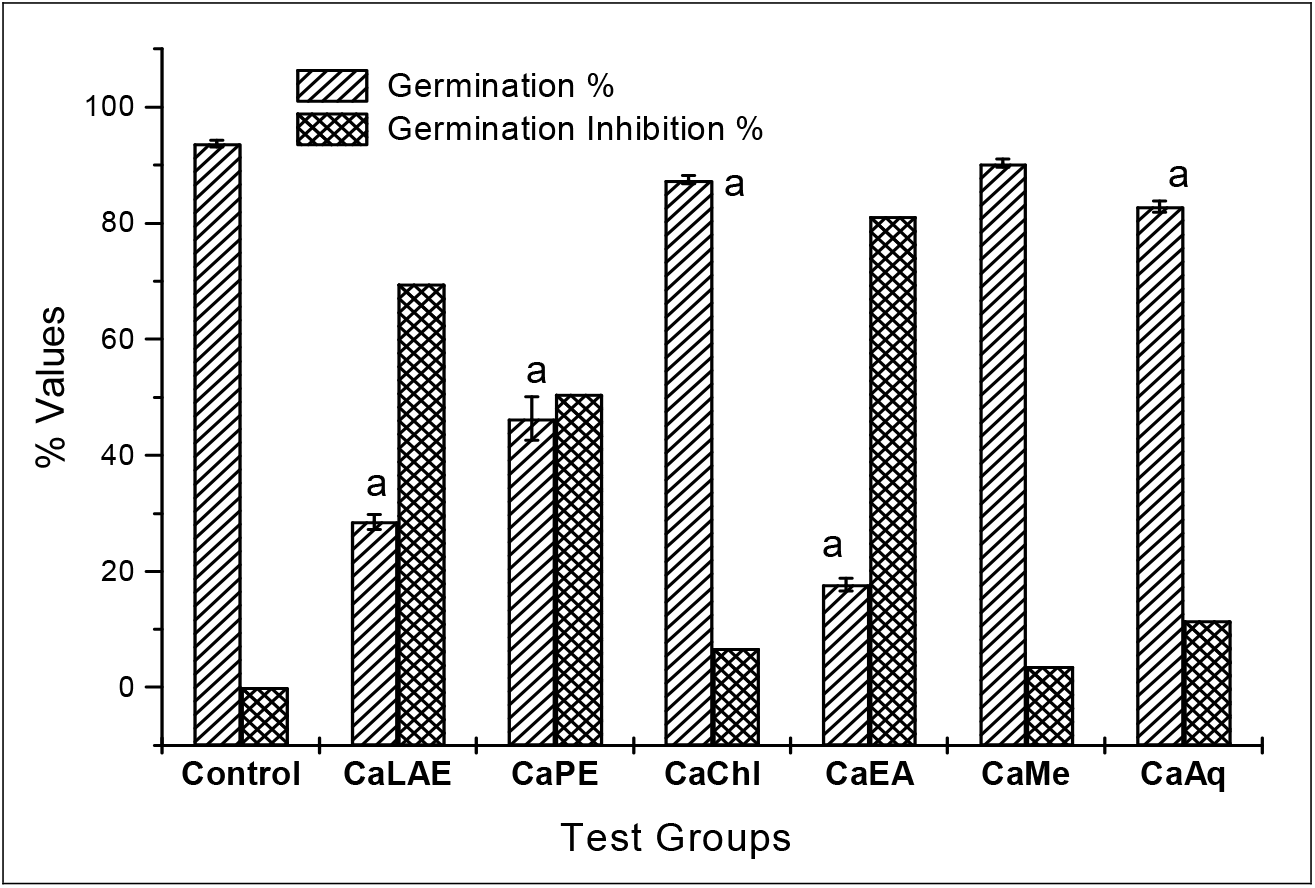
Mean germination % and germination inhibition % after 72 h of continuous treatment with 1 mg/mL concentration of different plant extracts. ^a^significant at *p<0.001*, ^b^at *p<0.01*, ^c^at *p<0.05* as compared to their respective control by Student’s t-test (two population); CaLAE-crude aqueous extract; CaPE-petroleum ether fraction; CaChl-chloroform fraction; CaEA-ethyl acetate fraction; CaMe-methanol fraction; and CaAq-aqueous fraction.

### 3.2. Growth retardation assay on sprouting monocot seedlings

All the six above mentioned extracts were tested for growth retardation effect on wheat seedlings, for up to 7 days of continuous treatment. After 7 days of incubation, the CaEA fraction was found to be the most effective growth retardant and CaAq as the least, both on root and shoot. The CaPE fraction also affected comparable to the CaEA. The CaEA stalled 99.7 and 99.5% root and shoot growth in respect to the control group (maintained with distilled water) at day 7 and found to be nine and four-fold more effective (growth inhibitory) than CaLAE and CaPE fractions on root cells and it reaches up to 122.6 and 1.25 fold on the shoot (Figure 02 and 04). They all significantly inhibited root growth at the level of 0.1% in every data recorded. If we consider dry weight accumulation, here also CaEA was the top inhibitor, 95.53% lesser dry matter accumulation was noted than the control; and CaPE was just 1% less effective than it (Figure 03). Whereas the CaAq fraction inhibited only 14% root and 15.2% shoot growth and 29.35% dry weight accumulation, which are least among all.

**Figure 02.**
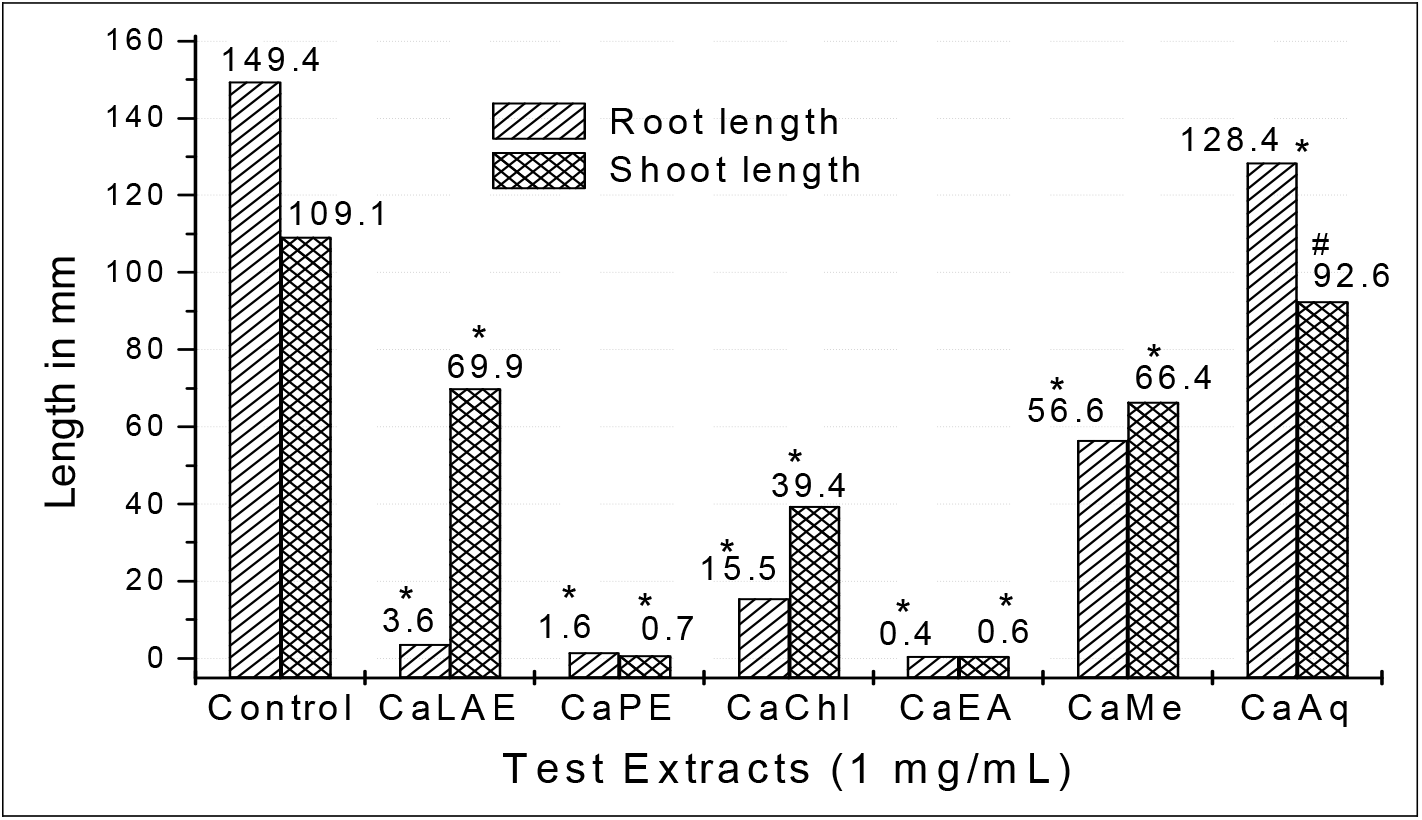
Effect of CaLAE and successive extract fractions of *C. asiaticum* leaves on wheat seedling growth on the 7^th^ day of treatment, *significant at *p<0.001*, ^@^at *p<0.01*, ^#^at *p<0.05* as compared to their respective control by Student’s t-test (two population); CaLAE-crude aqueous extract; CaPE-petroleum ether fraction; CaChl-chloroform fraction; CaEA-ethyl acetate fraction; CaMe-methanol fraction; and CaAq-aqueous fraction.

**Figure 03.**
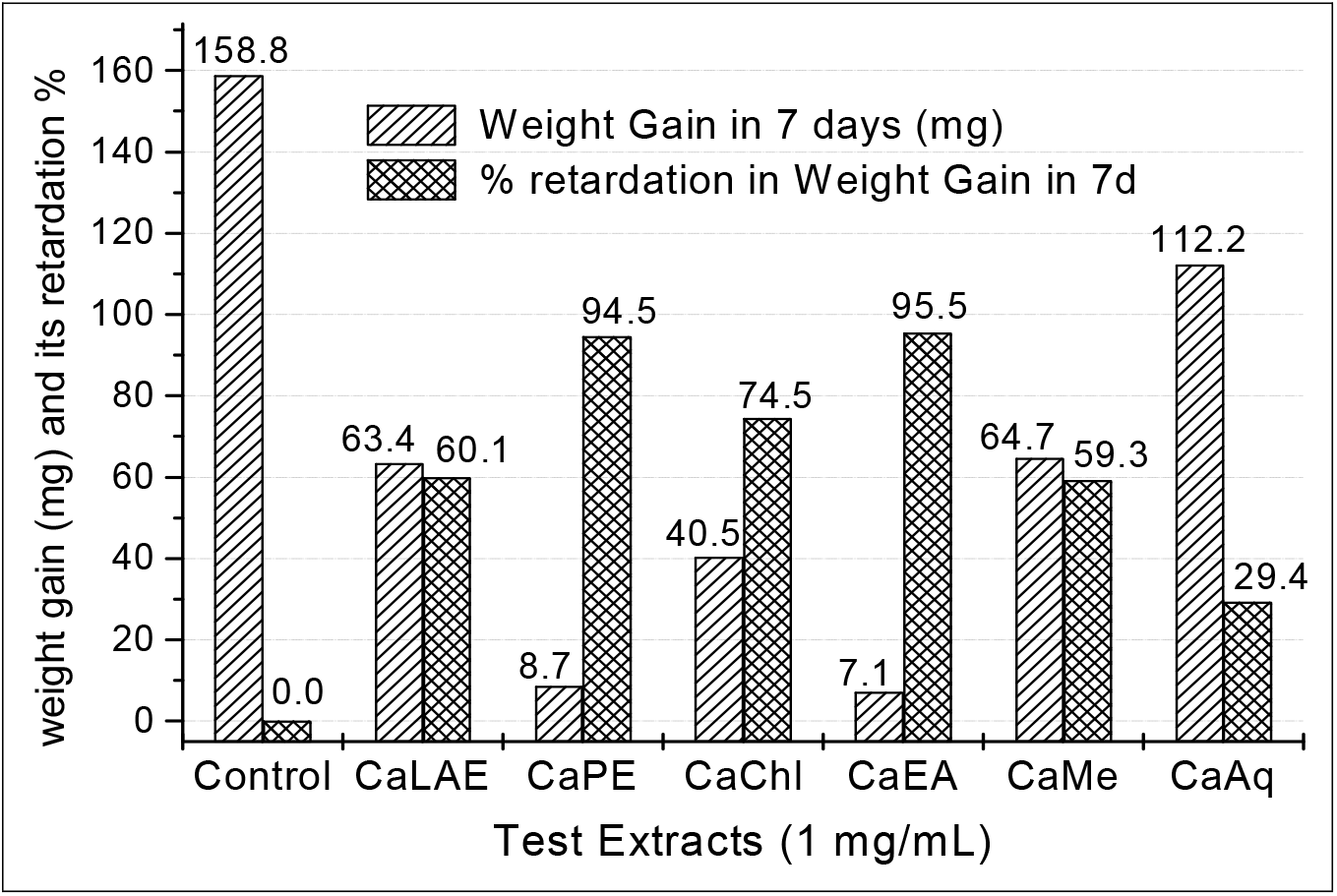
Effect of CaLAE and successive extract fractions of *C. asiaticum* leaves on dry weight (mg) accumulation of wheat seedlings, after 7 days of incubation. CaLAE-crude aqueous extract; CaPE-petroleum ether fraction; CaChl-chloroform fraction; CaEA-ethyl acetate fraction; CaMe-methanol fraction; and CaAq-aqueous fraction.

**Figure 04.**
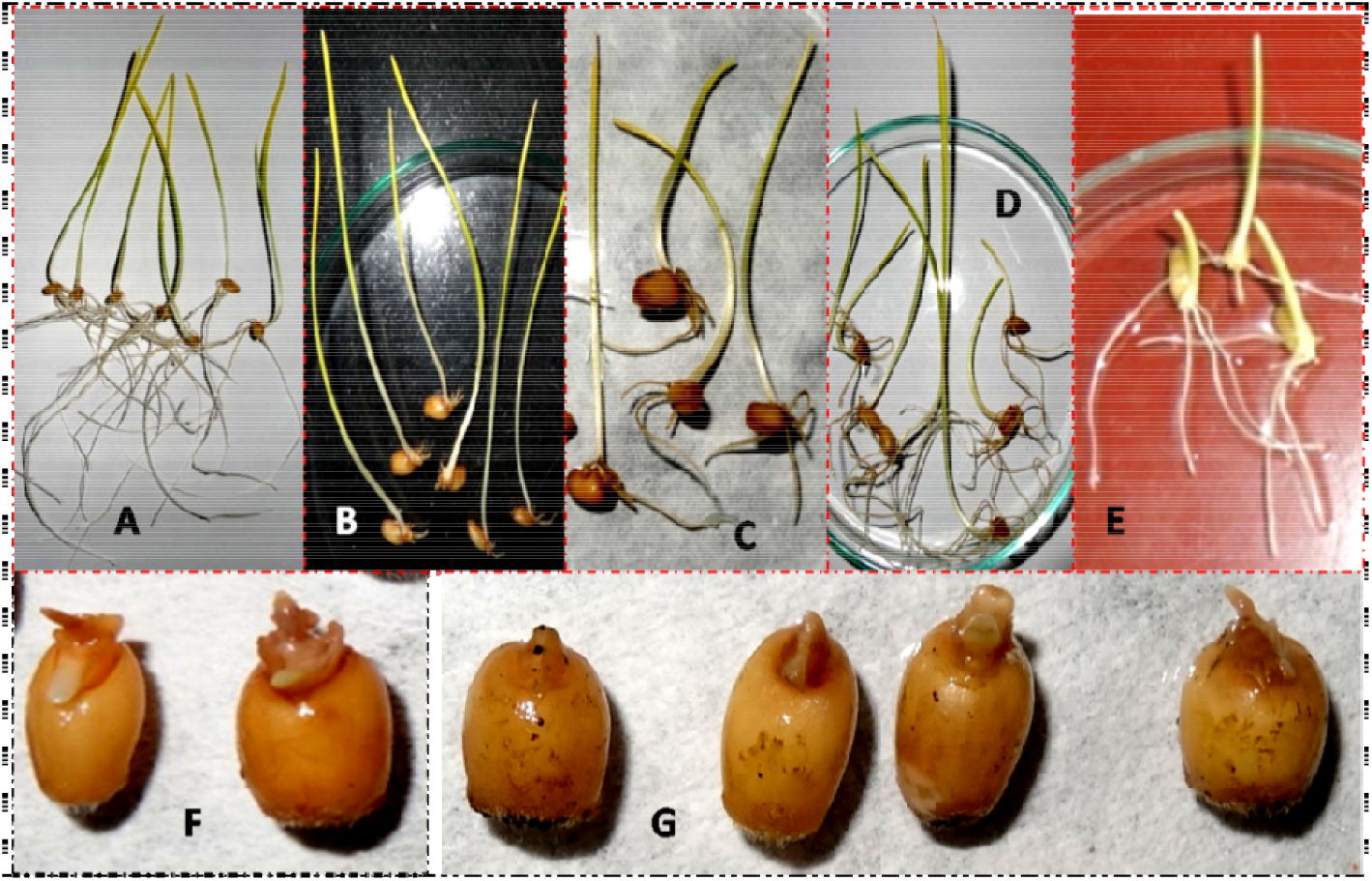
Showing the growth retardation effects of the CaLAE and successive extract fractions of *C. asiaticum* leaves on wheat seedlings, after 7 days of treatment. A-Control, B-crude aqueous extract (CaLAE), C-chloroform fraction (CaChl), D-methanol fraction (CaMe), E-aqueous fraction (CaAq), F-petroleum ether fraction (CaPE), G-ethyl acetate fraction (CaEA).

### 3.3. Growth retardation assay on sprouting dicot seedlings

We also used a different model i.e. chickpea (*Cicer arietinum*), a dicot seed, for growth retardation study. This study again highlights CaEA as the most effective among the six test extracts, with complete growth stall at day 6 in the lower concentration (Table 01). But remarkable in this model CaPE has a much lesser effectiveness (only 32.5% at 1 mg/mL) than the previous monocot model (98.95% at 1 mg/mL). Both CaLAE and CaEA blocked 100% root growth in 2 mg/mL concentration, at 48 h of incubation. With the lower concentration, these percentages were 96.7 and 100% respectively for CaLAE and CaEA treatment at the final hour.

**Table 01.** Pooled data showing chickpea root growth retardation effects of CaLAE and successive extract fractions of *C. asiaticum* leaves, up to 144 h

| Extract treatment | Dose (mg/mL) | ROOT LENGTH (mm) Mean±SEM |  |  |  |  | RGI at 144 h |
| --- | --- | --- | --- | --- | --- | --- | --- |
|  |  | 00 h | 24 h | 48 h | 72 h | 144 h |  |
| Control | 00 | 16.0±1.5 | 42.0±2.7 | 114.0±1.7 | 173.0±1.0 | 279.6±2.1 | 0.00 |
| CaLAE | 1 | 16.8±1.0 | 18.2±3.0<sup>a</sup> | 5.0±3.3<sup>a</sup> | 6.0±4.0<sup>a</sup> | 9.2±6.1<sup>a</sup> | **96.7** |
|  | 2 | 15.8±0.8 | 17.6±1.4<sup>a</sup> | 0.0±0.0<sup>a*</sup> | 0.0±0.0<sup>a*</sup> | 0.0±0.0<sup>a*</sup> | **100** |
| CaPE | 1 | 15.0±1.6 | 34.6± 7.7<sup>b</sup> | 81.4±4.4<sup>a</sup> | 116.6±6.7<sup>a</sup> | 188.6±3.7<sup>a</sup> | 32.5 |
|  | 2 | 14.6±1.1 | 20.2±3.6<sup>a</sup> | 47.0±0.9<sup>a</sup> | 72.6±3.1<sup>a</sup> | 128.4±1.7<sup>a</sup> | 54.1 |
| CaChl | 1 | 15.0±2.2 | 19.2±1.8<sup>a</sup> | 34.6±3.6<sup>a</sup> | 78.4±7.1<sup>a</sup> | 126.8±6.5<sup>a</sup> | 54.6 |
|  | 2 | 14.4±1.5 | 17.6±2.0<sup>a</sup> | 39.8±1.5<sup>a</sup> | 69.0±3.5<sup>a</sup> | 94.4±1.9<sup>a</sup> | 66.2 |
| CaEA | 1 | 14.6±1.4 | 17.2±1.2<sup>a</sup> | 19.2±0.5<sup>a</sup> | 9.2±3.8<sup>a</sup> | 0.00±0.0<sup>a*</sup> | **100** |
|  | 2 | 15.2±1.8 | 16.6±0.9<sup>a</sup> | 0.0±0.0<sup>a*</sup> | 0.0±0.0<sup>a*</sup> | 0.0±0.0<sup>a*</sup> | **100** |
| CaMe | 1 | 15.2±1.7 | 23.0±3.5<sup>a</sup> | 48.8±4.8<sup>a</sup> | 75.4±9.2<sup>a</sup> | 179.4±6.8<sup>a</sup> | 35.8 |
|  | 2 | 15.8±1.4 | 25.4±2.7<sup>a</sup> | 35.4±2.5<sup>a</sup> | 40.8±3.1<sup>a</sup> | 142.6±3.9<sup>a</sup> | 48.9 |
| CaAq | 1 | 16.2±0.8 | 27.6±2.1<sup>a</sup> | 72.8±2.9<sup>a</sup> | 130.4±3.6<sup>a</sup> | 198.6±3.4<sup>a</sup> | 28.9 |
|  | 2 | 15.6±1.5 | 25.8±4.3<sup>a</sup> | 42.8±4.0<sup>a</sup> | 61.0±7.0<sup>a</sup> | 140.2±4.7<sup>a</sup> | 49.8 |

<sup>a</sup> significant at $p<0.001$ , <sup>b</sup> at $p<0.01$ , <sup>c</sup> at $p<0.05$ as compared to their respective control by Student's t-test (two population); \*- completely rotten; h- Hours; Conc- Concentration; CaLAE- crude aqueous extract; CaPE- petroleum ether fraction; CaChl- chloroform fraction; CaEA- ethyl acetate fraction; CaMe- methanol fraction; and CaAq- aqueous fraction.

### 3.4. Chironomid larval mortality assay

Among the most sensitive population of first instars larvae, CaEA, CaPE, and CaLAE treated groups were mostly affected among all (Table 02). LC_50_ values at 24 h for those three extracts were calculated very low respectively as 0.87, 1.3 and 1.48 mg/mL (Figure 05). For CaAq and CaME this was respectively 4.2 and 10.7 times of the most toxic CaEA fraction. In CaEA and CaLAE treated group, 100% larval mortality was achieved only in 24 h with 2.0 mg/mL dose and in the lowest dose (0.5 mg/mL) it required upto 72 h. Also in CaPE treated group, complete larval mortality was achieved in 24 h but with dose doubling (4.0 mg/mL). CaAq and CaME found to be very less toxic with much higher LC_50_ values respectively as 3.62 and 9.32 mg/mL at 24 h. When we consider LT_50_ for 1 mg/mL dose it was nearly 23 h for CaEA and 47 h both in CaLAE and CaChl, whereas CaPE remains in between them with approximately 30 h to affect similarly (Figure 06). Alike LC_50_for CaAq and CaME, LT_50_ for 1 mg/mL again reaches highest among all, more than 57 h.

**Figure 05.**
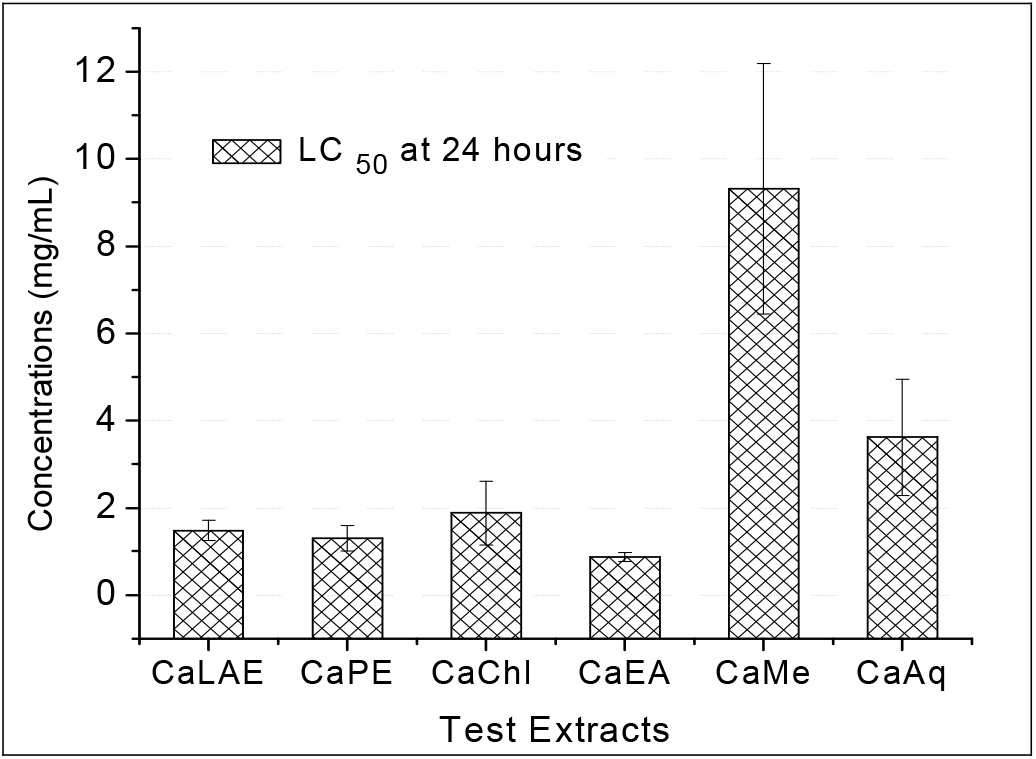
Showing the LC_50_ values of the different extract fractions of *C. asiaticum* leaves. CaLAE-crude aqueous extract; CaPE-petroleum ether fraction; CaChl-chloroform fraction; CaEA-ethyl acetate fraction; CaMe-methanol fraction; and CaAq-aqueous fraction.

**Figure 06.**
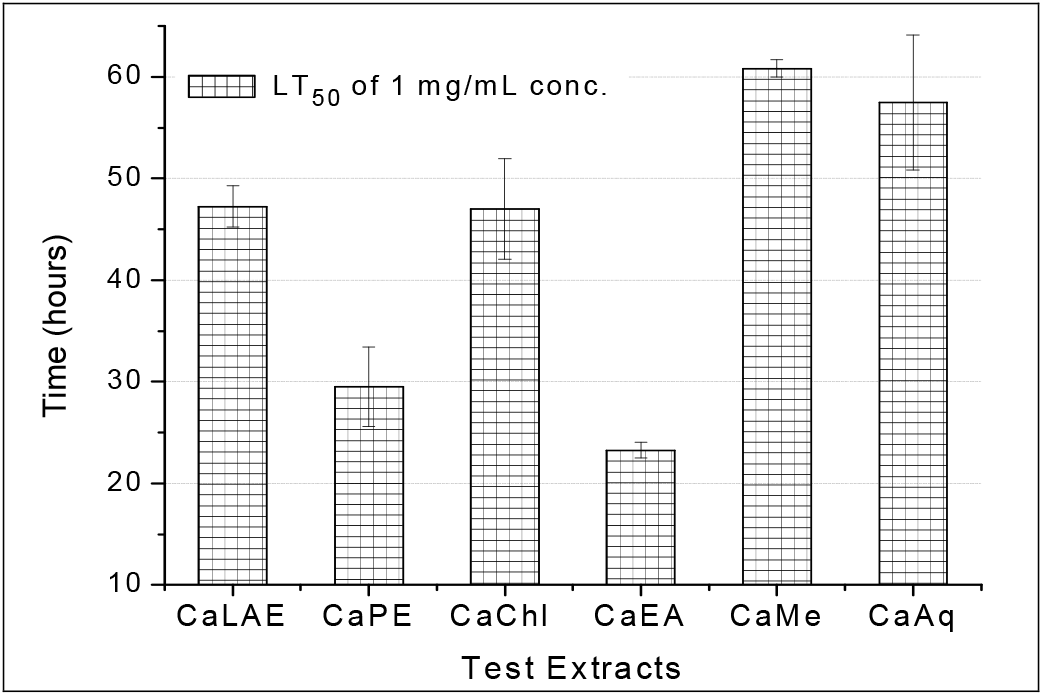
Showing the LT_50_ values of different extract fractions of *C. asiaticum* leaves having 1 mg/mL concentration. CaLAE-crude aqueous extract; CaPE-petroleum ether fraction; CaChl-chloroform fraction; CaEA-ethyl acetate fraction; CaMe-methanol fraction; and CaAq-aqueous fraction.

**Table 02.**
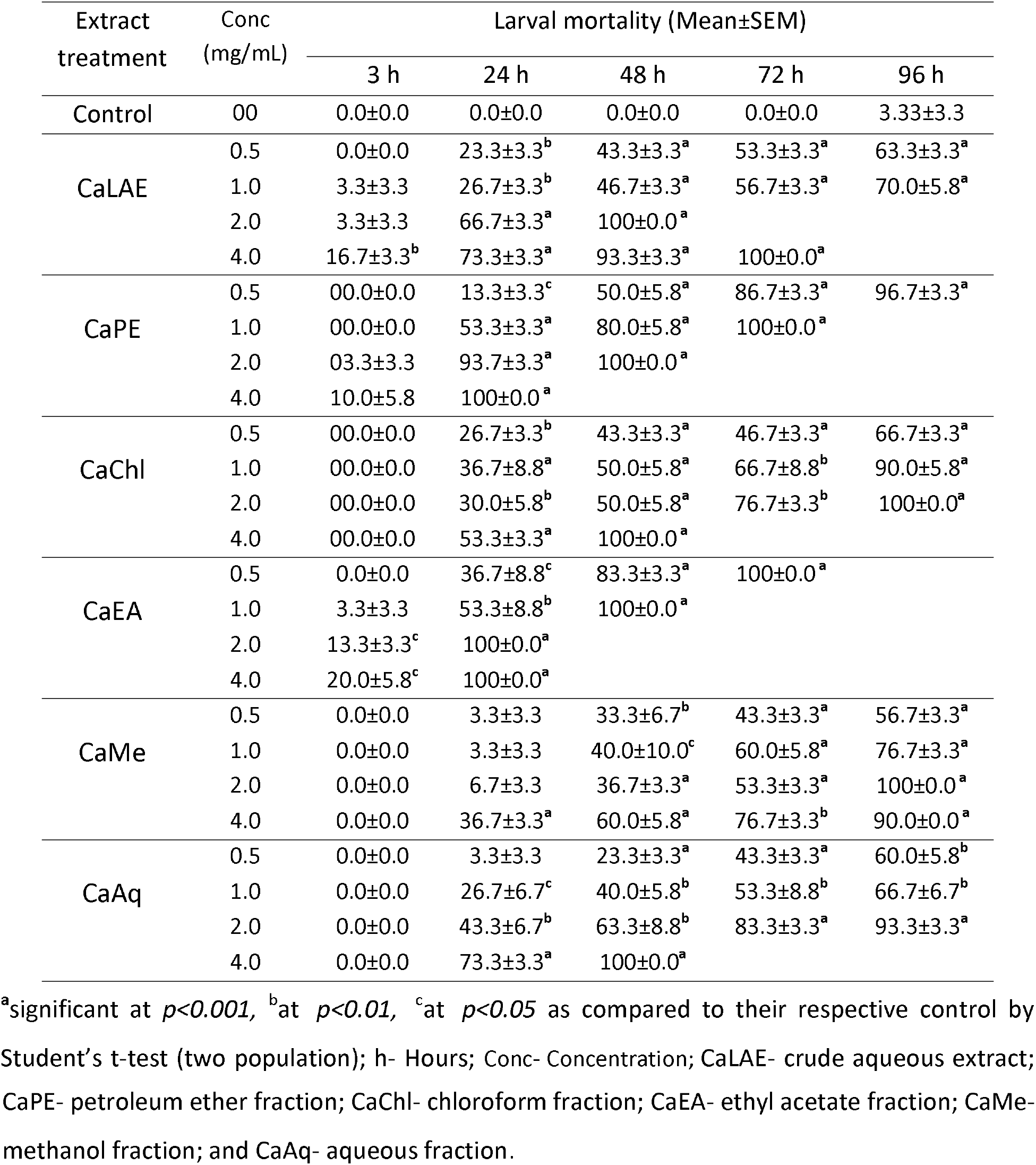
Pooled data showing dose and time-dependent effect of *C. asiaticum* leaves extract fractions on Chironomid larval mortality

CaPE-petroleum ether fraction; CaChl-chloroform fraction; CaEA-ethyl acetate fraction; CaMe-methanol fraction; and CaAq-aqueous fraction.

## 4. Discussions

Plant-plant interactions, mostly growth inhibitory allelopathic effects draw the major attention of the researchers in screening of cytotoxic phytochemicals due to their cost efficiency and prompt outcome. In continuation to our previous comparative antimicrobial and antioxidant study with the crude aqueous extract and five different polarity gradient solvent-mediated Soxhlet fractions of *Crinum asiaticum* leaves (Goswami et al., 2020), the same extract fractions were further assayed for their comparative phyotoxic effects. They were tested on wheat seedlings for germination inhibition and growth retardation effects and the ethyl acetate fraction (CaEA) was found as the most potent germination inhibitor, followed by the crude aqueous (CaLAE) and petroleum ether (CaPE) fractions, and the remaining three i.e. CaChl, CaMe, and CaAq induced very lesser effect. As an antioxidant rich plant exert several health benefits, precautions can be taken against free radical mediated damages by eating ample amount of antioxidant-rich foodstuffs. It is well established fact that plant phenolics and flavonoids have strong antioxidant activities and thus they are able to prevent certain degenerative diseases and also reduces the risk of cardiovascular disease as well as certain cancers (Sun *et al*., 2002).

To evaluate the bioactivity of phytochemicals diverse experimental models, such as cell cultures, membrane systems, different plants or animal model systems, and finally clinical trials are used. Their preliminary antiproliferative and cyto-genotoxic effects were determined using plant-based models, by simple germination and growth inhibition assays. Exact compositions of the potent plant extract(s) are usually required for pharmaceutical uses. In plants these compounds maintain dormancy of specific organ(s) or of the whole plant, are produced in any or particular plant part and may exert dose-dependent effect to other organisms and also dependent on the solvent for proper extraction. Dose optimization is also crucial and many trials required getting the desired outcome. The preliminary evaluation of phytotoxic effects of such chemical can be performed using plant models to ensure cost-effective, rapid and easy to handle setups (Angayarkanni *et al*., 2007; Camparoto *et al*., 2002; Fachinetto *et al*., 2007; Fiskesjo, 1985). A previous study indicates, alteration in microtubules or their associated proteins leads to stall root elongation of wheat seedlings (Armbruster *et al*., 1991). Stunted plant growth indicates retardation in cell division by antiproliferative and cytotoxic effects (Yildiz *et al*., 2009). As reported, the latex of *Calotropsis procera* both in crude and methanolic extraction induced antimitotic activity in *A. cepa* (Sehgal *et al*., 2006). These test systems have been accepted by various scientists because of resulting similarly, both on animals and plants (Teixeira *et al*., 2003; Vicentini *et al*., 2001). Another plant model used in our study, the chickpea seedlings, also found to be reactive to exogenous plant extract exposures. According to Carita and Marin-Morales, 2008, plant models are the efficient way for assessment of allelopathic genotoxicity study, through chromosomal anomalies inducing potentials. Therefore, the use of these model systems is cheap, simple and easy to handle for cytogenotoxicity evaluation. In addition, these data are very much correlated to the experimental records using animal models.

From the comparative monocot seedling growth retardation assay, after seven days of continuous treatment, the CaEA and CaAq fractions were found to be the most and least effective respectively, on both root and shoot growth. The CaPE also affected somehow parallel to CaEA; and if we consider meristematic growth as the result of cell division then, CaEA found to be 9 and 4 fold more effective than CaLAE and CaPE on root tip cells and 122.6 and 1.25 fold on the shoot, in respect to cell proliferation inhibition. Additionally, roots show higher sensitivity to the extracts than the shoots and that may be due to the fact that roots were given direct exposures of the extracts but shoots show any effects only if the active principle(s) are absorbed and circulated. According to Fiskesjo (1985), root growth retardation is the outcome of the suppression of cell divisions and incorporation of chromosomal aberrations (Fiskesjo, 1985). Narciclasine 1, an alkaloid with potent antitumor activity, was isolated from *Narcissus sp*. bulbs, primarily based on the report of its inhibitory effects on the growth of wheat grains (Ceriotti, 1967). *Narcissus sp*. is also a member of the Amaryllidaceae family of which the plant *C. asiaticum* belongs.

The growth retardation assay of sprouting dicot (chickpea-*C. arietinum*) seedling also highlights the same fraction (CaEA) as the most effective amongst all. But remarkable in this model CaPE has much lesser effect than the monocot model. Both CaEA and CaLAE blocked 100 % root growth in 2 mg/mL concentration, at 48 h. Thus CaEA and CaLAE were selected for further dose and time dependent-comparative studies and on basis of the comparative assay for their effect on monocot growth, CaEA found to be approximately 1.4 times more effective than CaLAE in respect to their IC_50_ (inhibitory concentration where 50% growth or germination inhibited) for germination inhibition and root growth study (data not shown here).

A widely used toxicity bioassay, the Chironomid larvae mortality assay, also established CaEA, CaPE, and CaLAE as more toxic than the other fractions. A larval toxicity assay is considered to be valid only if, less than 20 % of the larvae show immobilization or other signs of stress (unusual behavior like trapping at the water surface, discoloration, etc.) at the end of the test in control set (Weltje et al., 2010). First instars larvae are of higher sensitivity than the older, as EC_5_o (effective concentration where 50% of the individuals are dead or immobile) values could differ highly among the first and fourth instars of *C. riparius*, for e.g. up to 950 times for Cadmium (Williams et al., 1986). Among the most sensitive population of first instars larvae of Chironomid, considering LC_50_ values for 24 h and LT_50_ for 1 mg/mL dose the ethyl acetate fraction found to be most toxic, followed by CaPE and CaLAE. For CaAq and CaME, LC_50_ values were respectively 4.2 and 10.7 times and LT_50_ values (for 1 mg/mL) for both near about 2.5 times of the most toxic CaEA fraction, which establishes them as far-less toxic fractions.

We carried out our initial studies using plant root apical meristem cells, which is actually an *in vivo* model. Cytotoxicity analyses, using such *in vivo* plant systems, are validated worldwide by various researchers who did this together with *in vitro* animal systems and reported similar outcomes (Fachinetto *et al*., 2007; Fiskesjo, 1994; Vicentini *et al*., 2001). Root growth stunted if the cell cycle kinetics is disrupted, which may due to of several phenomenons like cellular accumulation at interphase, and/or by disturbing the normal process of the mitotic spindle formation, and/or by inducing chromosomal anomalies leading to a reduction in the mitotic indexes (Yadav, 1986; Vyuyan, 2002).

The plant *Terminalia citrina* (Gaertn.) Roxb., a traditional medicinal plant of the Combretaceae family, which was previously reported for antimicrobial tannins from its fruit, is now recently in 2018 reported again by Muhit et al., for inhibiting estradiol (E2) induced proliferation of human breast cancer cell lines (MCF-7 and T47D). Several plants with antioxidant properties and higher phenolics content have been widely reported for their ability to reduce the risk and slowdown the rate of progression of several types of cancer diseases mainly through the prevention of cellular oxidation (Pereira do Amaral et al., 2012). The CaEA fraction, found to be the most hazardous here, was previously reported to have significantly higher quantity of phenolics and flavonoids as its phytochemistiry, and with distinct antimicrobial properties (Goswami et al., 2020). Thus may be a promising resource in future drug development and therapeutics to treat deadly diseases, such as cancer. On the other hand, the most nontoxic fraction i.e. the last Soxhlet mediated aqueous fraction (CaAq), which has also been reported to have the highest antimicrobial properties among all the tested extracts (Goswami et al., 2020) might be used in live stock maintenance in spite of chemical antibiotics with long-term side effects on us.

## 5. Conclusions

The crude aqueous, as well as the non polar extract fractions made up of petroleum ether and ethyl acetate is directly causing significant toxic effects on both meristematic tissue and aquatic midges. The phytochemistry behind these toxicities may reason the repulsion of insects and herbivores from the plant. The last aqueous fraction of the successive extraction was found to have minimal phytotoxic and lethal effects and, as reported earlier, have potent antioxidant and antibacterial activities. Thus the present study indicates antimicrobial and antioxidant prospects with minimum toxicity of the last aqueous extract fraction of *Crinum asiaticum* leaves and its future beneficial use in livestock maintenance. Further, it is needed to explore anticancer pharmaceutical prospects of the ethyl acetate extract fraction.

## List of abbreviations

CaLAE: Crude **l**eaf **a**queous **e**xtract of ***C**. **a**siaticum*
CaPE: Successive **P**etroleum **e**ther fraction of ***C**. **a**siaticum*
CaChl: Successive **Chl**oroform fraction of ***C**. **a**siaticum*
CaEA: Successive **E**thyl **a**cetate fraction of ***C**. **a**siaticum*
CaMe: Successive **Me**thanol fraction of ***C**. **a**siaticum*
CaAq: Successive **Aq**ueous fraction of ***C**. **a**siaticum*

## Acknowledgement

The authors gratefully acknowledge Prof. Ambarish Mukherjee, Department of Botany, B.U. for authentication of of the plant species and also acknowledge the financial support of the UGC {F.No.42-563/2013 (SR) dt. 22.3.13}, and infrastructural supports of DST-PURSE, DST-FIST, UGC-DRS to the Department of Zoology, The University of Burdwan, West Bengal, India.

