## Supplemental Table and figures for "Comparative phytotoxicity and chironomid toxicity assessments of leaf successive extract fractions of Asiatic poison bulb, *Crinum asiaticum* L"

*SUPPLIMENTARY*

The extraction procedure through Soxhlet is described schematically, in Flow chart below.

**3.1.1.** **Screening and selection of the most active fraction of *C. asiaticum*: Germination inhibition assay on wheat**

**Table 01.** Effect of CaLAE and other extract fractions of *C. asiaticum* leaves*,* on germination of wheat seeds, after 72 h of treatment

| Extract treatment (1mg/mL) | Germination % (72 h) | | GI % |
| --- | --- | --- | --- |
|  | Range | Mean±SEM |  |
| Control | 91.4-94.3 | 93.7±0.6 | 0.00 |
| CaLAE | 25.7-31.4 | 28.6±1.3^a^ | 69.5 |
| CaPE | 31.4-51.4 | 46.3± 3.8^a^ | 50.6 |
| CaChl | 85.7-88.6 | 87.4±0.7^a^ | 06.7 |
| CaEA | 14.3-20.0 | 17.7±1.1^a^ | 81.1 |
| CaMe | 88.6-91.4 | 90.3±0.7 | 3.66 |
| CaAq | 80.0-85.7 | 82.9±0.9^a^ | 11.6 |

^a^ significant at *p<0.001,* ^b^ at *p<0.01,* ^c^ at *p<0.05* as compared to their respective control by Student’s t-test (two population); h- Hours; GI- Germination Inhibition in comparison to control group; Conc– concentration;

**3.1.2.** **Growth retardation assay on sprouting monocot seedling by CaLAE and successive soxhlet fractions of *C. asiaticum***

**Table 02.** Pooled data showing the influence of the successive extracts of *C. asiaticum* on the root growth of wheat seedlings, up to day seven

| Extract treatment (1mg/mL) | ROOT LENGTH (mm) | | | | | | |
| --- | --- | --- | --- | --- | --- | --- | --- |
|  | **24 h** | | **4^th^ day (96 h)** | | **7^th^ day (168 h)** | | |
|  | RANGE | Mean±SEM | RANGE | Mean±SEM | RANGE | Mean±SEM | RGI% |
| Control | 11-20 | 15.6±0.6 | 47-68 | 58.8±1.3 | 136-162 | 149.4±1.8 | 0.00 |
| CaLAE | 00-02 | 1.2±0.1^a^ | 01-03 | 1.4±0.1^a^ | 02-06 | 3.6±0.3^a^ | **97.59** |
| CaPE | 00-01 | 0.6±0.2^a^ | 00-02 | 1.2±0.1^a^ | 00-04 | 1.6±0.3^a^ | 98.95 |
| CaChl | 01-08 | 3.3±0.5^a^ | 05-13 | 7.5±0.5^a^ | 09-22 | 15.5±0.9^a^ | 89.64 |
| CaEA | 00-01 | 0.14±0.1^a^ | 00-01 | 0.6±0.2^a^ | 00-01 | 0.4±0.2^a^ | **99.72** |
| CaMe | 06-13 | 9.3±0.5^a^ | 12-37 | 24.7±1.2^a^ | 41-72 | 56.6±2.0^a^ | 62.10 |
| CaAq | 05-17 | 9.86±0.9^a^ | 31-50 | 39.67±1.4^a^ | 111-141 | 128.38±1.6^a^ | 14.06 |

^a^ significant at *p<0.001,* ^b^ at *p<0.01,* ^c^ at *p<0.05* as compared to their respective control by Student’s t-test (two population); h- Hours; RGI-Root Growth Inhibition.

**Table 03.** Pooled data showing the influence of the successive extracts of *C. asiaticum* on the shoot growth of wheat seedlings, up to day seven

| Extract treatment (1mg/mL) | SOOT LENGTH (mm) | | | | | | |
| --- | --- | --- | --- | --- | --- | --- | --- |
|  | **24 h** | | **4^th^ day (96 h)** | | **7^th^ day (168 h)** | | |
|  | RANGE | Mean±SEM | RANGE | Mean±SEM | RANGE | Mean±SEM | SGI% |
| Control | 09-12 | 10.6±0.4 | 42-52 | 47.0±1.4 | 95-125 | 109.1±4.1 | 0.00 |
| CaLAE | 04-08 | 5.6±0.6^a^ | 25-32 | 28.4±0.9^a^ | 64-75 | 69.9±1.3^a^ | 35.99 |
| CaPE | 00-01 | 0.6±0.2^a^ | 01-01 | 1.0±0.0^a^ | 00-01 | 0.7±0.2^a^ | **99.35** |
| CaChl | 01-03 | 1.7±0.3^a^ | 12-22 | 16.6±1.6^a^ | 28-52 | 39.4±3.6^a^ | 63.87 |
| CaEA | 00-01 | 0.6±0.2^a^ | 00-01 | 0.6±0.2^a^ | 00-02 | 0.6±0.29^a^ | **99.48** |
| CaMe | 02-05 | 3.9±0.4^a^ | 14-31 | 21.9±2.3^a^ | 42-110 | 66.4±8.4^a^ | 39.13 |
| CaAq | 08-11 | 9.6±0.4 | 28-44 | 33.9±2.3^a^ | 73-112 | 92.6±5.2^c^ | 15.18 |

^a^ significant at *p<0.001,* ^b^ at *p<0.01,* ^c^ at *p<0.05* as compared to their respective control by Student’s t-test (two population); h- Hours; SGI- Shoot Growth Inhibition.

**Table 04.** Pooled data showing the influence of CaLAE and the successive extracts of *C. asiaticum* leaves on root-shoot growth of wheat seedlings on day seven

| Extract treatment (1mg/mL) | ROOT LENGTH (mm) | | | SOOT LENGTH (mm) | | |
| --- | --- | --- | --- | --- | --- | --- |
|  | RANGE | Mean±SEM | RGI % | RANGE | Mean±SEM | SGI % |
| Control | 136-162 | 149.4±1.8 | 00.00 | 95-125 | 109.1±4.1 | 00.00 |
| CaLAE | 02-06 | 3.6±0.3^a^ | 97.59 | 64-75 | 69.9±1.3^a^ | 35.99 |
| CaPE | 00-04 | 1.6±0.3^a^ | 98.95 | 00-01 | 0.7±0.2^a^ | 99.35 |
| CaChl | 09-22 | 15.5±0.9^a^ | 89.64 | 28-52 | 39.4±3.6^a^ | 63.87 |
| CaEA | 00-01 | 0.4±0.2^a^ | 99.72 | 00-02 | 0.6±0.29^a^ | 99.48 |
| CaMe | 41-72 | 56.6±2.0^a^ | 62.10 | 42-110 | 66.4±8.4^a^ | 39.13 |
| CaAq | 111-141 | 128.4±1.6^a^ | 14.06 | 73-112 | 92.6±5.2^c^ | 15.18 |

^a^ significant at *p<0.001,* ^b^ at *p<0.01,* ^c^ at *p<0.05* as compared to their respective control by Student’s t-test (two population); h- Hours; SGI- Shoot Growth Inhibition.

**Table 05.** Influence of CaLAE and the successive extracts of *C. asiaticum* leaves on weight gain of wheat seedlings on day seven

| Extract treatment (1 mg/mL) | Weight gain in 7 d (mg) | % retardation in weight gain in 7d |
| --- | --- | --- |
| Control | 158.8 | 00.00 |
| CaLAE | 63.4 | 60.08 |
| CaPE | 08.7 | 94.52 |
| CaChl | 40.5 | 74.50 |
| CaEA | 07.1 | **95.53** |
| CaMe | 64.7 | 59.26 |
| CaAq | 112.2 | 29.35 |

**Comparative study of CaLAE and CaEA for their effect on monocot seedling germination and growth**

Cont A

31.25

62.5

125

250

500

Cont B

31.25

62.5

125

250

500

0

10

20

30

40

50

60

70

80

90

100

% Values

CaLAE	CaEA

Test Extract

Germination Inhibition %

Root Growth Inhibition %

**Figure 05.** Showing CaLAE and CaEA induced dose-dependent germination and growth retardation effects on wheat seedlings, at 72 h of incubation.

**Comparative study of the TLC fractions of CaEA for their effect on monocot seedling growth**

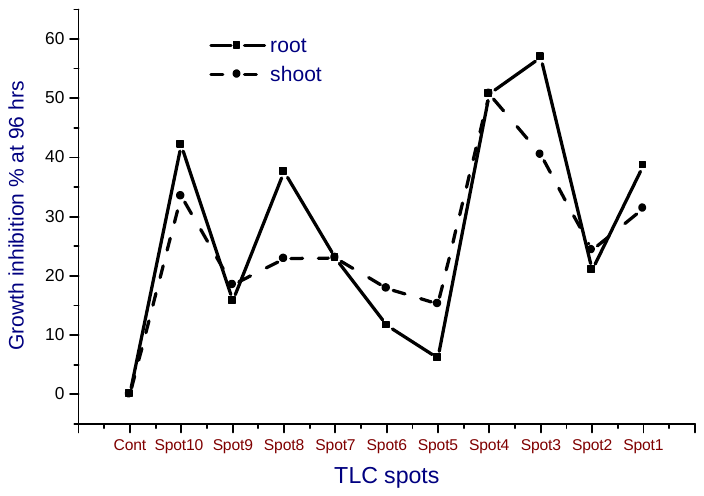

**Figure III.09.** Showing relative root-shoot growth retardation effects of TLC fractions of CaEA on wheat seedlings, at 96 h.

***4.4. Chironomid larval mortality assay***

**Table 03.** Showing LC_50_ (at 24 h) and LT_50_ (for 1 mg/mL) of the different extract fractions

of *C. asiaticum* leaves

| Extract treatment | LC_50_ for 24 h | LT_50_ of 1 mg/mL concentration |
| --- | --- | --- |
| CaLAE | 1.48±0.23 | 47.26±2.07 |
| CaPE | 1.3±0.29 | 29.5±3.9 |
| CaChl | 1.88±0.74 | 47.0±4.95 |
| CaEA | **0.87±0.1** | **23.26±0.75** |
| CaMe | 9.32±2.87 | 60.86±0.86 |
| CaAq | 3.62±1.33 | 57.49±6.64 |
